# Genetic interaction approaches reveal emerging roles of innexins in development : Insights from a novel *pannier–innexin-2* interaction during *Drosophila* embryogenesis

**DOI:** 10.64898/2026.06.22.733794

**Authors:** Spraha Bhandari, Franka Eckardt, Reinhard Bauer

## Abstract

Effective communication between cells is essential for the typical development and behaviour of an organism. In this context, gap junctions represent the most universally preserved components at cellular membranes of multicellular organisms, facilitating metabolic and electrical connections between cells. Disruptions in these junctions have been linked to various developmental abnormalities and pathological conditions in humans. The invertebrate gap junction proteins, referred to as innexins, exhibit conserved cellular and molecular mechanisms of functioning with their vertebrate counterparts, known as connexins. Consequently, they provide valuable means for studying and understanding the functions of gap junctions in development. In the *Drosophila* embryo, innexin-2 is expressed in the amnioserosa and ectoderm, where it is required for epithelial morphogenesis. Genetic depletion of *innexin-2* results in cuticular defects and embryonic lethality. Pannier, a GATA family transcription factor, is a key regulator of dorsal tissue development in *Drosophila* and is expressed in the amnioserosa, dorsal ectoderm and the dorsal vessel during embryogenesis. *Pannier* mutants exhibit defects in dorsal closure, cuticle formation, and cardiac specification. Although substantial evidence from vertebrate systems indicate that connexin expression is regulated by transcription factors such as GATA4, Nkx2.5, Tbx2, Tbx3, and Tbx5, whether a similar regulatory relationship exists between these transcription factors and gap junction proteins in *Drosophila* remains unknown. In this study, we investigate how *innexin* mediated intercellular communication impacts *pannier* dependent morphogenetic processes during *Drosophila* embryogenesis.

## Introduction

During *Drosophila* embryogenesis, intercellular communication via gap junctions is crucial for embryonic body wall epithelia development, nervous system development, proper dorsal closure and formation of internal organs (Bauer *et al*., 2004; Lehmann *et al*., 2006; Ostrowski *et al*., 2008; Giuliani *et al*., 2013; Bauer *et al*., 2002). The established knowledge now encompasses the mRNA expression levels of all and the protein expression profiles of certain gap junction proteins, called innexins, throughout the different embryonic stages (Stebbings *et al*., 2002; Bauer *et al*., 2004; Lehmann *et al*., 2006). Among the eight *innexin* genes identified in *Drosophila, innexin-2* is the most widely expressed, being present across multiple tissues during development. Innexin-2 forms both the homomeric and heteromeric channels, notably interacting via its C-terminal domain with innexin-3 to form heteromeric complexes (Lehmann *et al*., 2006).

Functional studies demonstrate that appropriate levels of innexin-2 are required for normal body wall epithelial morphogenesis in fly embryos (Bauer *et al*., 2004). Interestingly, large-scale mutagenesis screens have also identified several key developmental regulators involved in embryonic development, including *pannier*, a GATA family transcription factor expressed in the ectoderm and amnioserosa. *Pannier* plays a crucial role in dorso-medial region specification and the regulation of dorsal closure during embryogenesis (Wieschaus & Volhard, 2016; Herranz & Morata, 2001). Homozygous nulls (*pnr*^*VX6-/-*^) of *pannier* fail to complete dorsal closure, resulting in lethality (Herranz & Morata, 2001; Heitzler *et al*, 1996). We examined innexin function during embryogenesis by analysing genetic interactions between *innexin-2* and *pannier*. The overlapping spatio-temporal expression patterns of *pannier* and *innexins*, along with their roles in embryonic development, the lethality observed in their respective loss-of-function mutants, and their participation in common developmental signalling pathways, led us to explore potential genetic interactions between the two genes using the *Drosophila* embryo as our model system (Figure 1). Towards the same, we carried out the genetic knockdown of *innexin-2*, using the epithelia specific *69B-Gal4* driver, in a *pannier* heterozygous null background. Experimental embryos (*UAS-inx2RNAi/+; pnr*^*VX6*^*/69B-Gal4*) exhibited marked defects in embryo size, shape, and body wall epithelial organisation, revealing a previously unrecognised link between gap junction components and transcriptional regulation during embryonic epithelial morphogenesis. In vertebrates, gap junction proteins known as connexins are regulated in a cell- and tissue-specific manner by distinct transcription factors. It is well established that GATA-4, a vertebrate homolog of pannier, binds to the promoter of Cx40 in both developing and mature atrial myocytes (Linhares *et al*., 2004). Furthermore, recent studies demonstrate that the synergistic overexpression of GATA-4 and MEF2C in human umbilical cord mesenchymal stem cells enhances the expression of cardiac genes and proteins, thereby influencing cell fate decisions during embryonic development (Razzaq *et al*., 2022).

**Figure 1:**
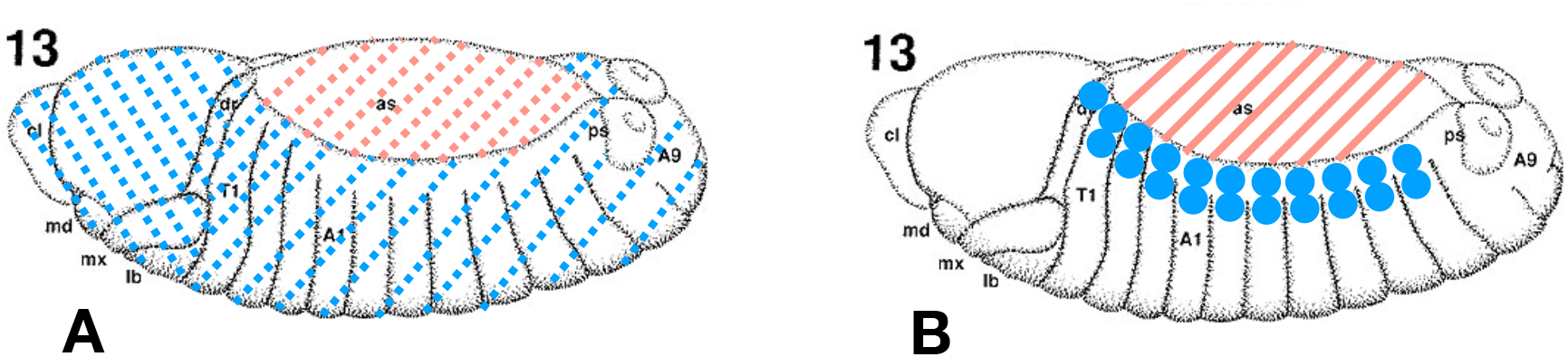
Schematic illustration of expression domains of innexin-2 (A) and pannier (B) in the ectoderm and amnioserosa of stage 13 *Drosophila* embryos. Pink indicates the amnioserosa, while blue marks the ectoderm.

## Materials and methods

### Fly culture and stocks

All *Drosophila* strains were cultured and maintained in regular corn flour media at 25°C. *Canton-S* was used as the wild-type control strain in our experiments. The *pannier* null mutant line, *pnr*^*VX6*^*/TM6B, Tb, Hu, e*, utilised in this study, was kindly provided by Marc Haenlin (Center for Integrative Biology, University of Toulouse, CNRS, UPS, Toulouse, France). The *UASwizInx2* line employed for RNAi mediated knockdown experiments was from Reinhard Bauer (LIMES Institute, University of Bonn, Bonn, Germany). The *69B-Gal4* and *pnr-Gal4* driver lines were procured from the Bloomington Drosophila Stock Center (BDSC 1774 and BDSC 3039, respectively). To visualise *pnr-Gal4* expression in embryos, a *UAS-10X-RFP* reporter line was used. The genotypes of all fly lines used in this study are listed in Supplementary Table 1.

### Embryo harvesting, staging and processing

Eggs from different fly lines were collected on yeasted apple juice plates for 2 hrs at 25°C in collection cages, followed by an additional 9 hrs and 12 hrs incubation at the same temperature to allow development up to the respective stages. The staged embryos were then collected using a nylon mesh strainer, dechorionated in 4% sodium hypochlorite (bleach) for 10 min, and washed three times with milliQ water. Fixation was subsequently carried out in a 1:1 mixture of heptane and 4% paraformaldehyde (PFA) with vigorous shaking (approximately 275–300 rpm) for 20 min at 25°C. Following fixation, embryos accumulated at the interface between the two phases. The lower aqueous phase was removed and replaced with an equal volume of methanol, after which the embryos were vortexed for 1–2 min at room temperature to facilitate devitellinisation. Subsequently, the upper phase containing debris was discarded, and the remaining embryos were stored in methanol at −20°C until further use.

### Immunohistochemistry and confocal experiments

For immunostaining, fixed embryos were rehydrated through a graded series of methanol and 1X PBS washes (10 min each). This was followed by a 10 min wash in 0.1% 1X PTX (PBS with 0.1% Tween-20). Embryos were then blocked in 5% normal goat serum (NGS), prepared in 0.1% 1X PTX, for 2 hrs at room temperature to prevent non-specific antibody binding. Following the blocking step, embryos were incubated overnight at 4°C with primary antibodies prepared in 0.1% 1X PTX supplemented with 5% NGS. The following primary antibodies were used : Rabbit Anti-Innexin-2 (1:50, kind gift from Alexander Borst, MPI for Biological Intelligence, Martinsried, Germany), Guinea Pig Anti-coracle (1:600, kind gift from Richard Fehon, University of Chicago, U.S.A), Mouse Anti-spectrin (1:10, Developmental Studies Hybridoma Bank), Mouse Anti-RFP (1:600, Developmental Studies Hybridoma Bank). After primary antibody incubation, embryos were washed for 10 min in 0.1% 1X PTX before addition of secondary antibodies. Fluorophore conjugated secondary antibodies-Alexa Fluor 488, Alexa Fluor 568 and Alexa Fluor 647, were used at a dilution of 1:700, and prepared in 0.1% 1X PTX supplemented with 2% NGS to detect the respective primary antibodies. Embryos were incubated with the secondary antibodies for 4 hrs at room temperature before mounting. Immunostained embryos were mounted on frosted microscope slides using VECTASHIELD antifade mounting medium and covered with 22 × 40 mm glass coverslips. Imaging was performed using an FV3000 upright confocal microscope at NCBS confocal imaging facility. The multi-channel Z-stack images were acquired using 20X air and 60X oil immersion objectives, with a step size of 0.31µm. For Supplementary Figure 1, images were taken with a Zeiss LSM710 confocal microscope at 20x magnification at LIMES, Bonn, Germany.

### Genetic knockdown experiments

To achieve genetic knockdown of *innexin-2* in both the dorsal ectoderm and amnioserosa, the males of *UASwizInx2* fly line were crossed with the virgins of *pnr-Gal4/TM6B, Dfd-EYFP, Sb* driver line. Embryos lacking GFP expression were selected for immunostainings. To induce *innexin-2* knockdown across a broader epithelial domain, the *69B-Gal4* driver line was employed.

### Generation of *pnr*^*VX6-/-*^ homozygous null embryos

The fly lines *pnr*^*VX6*^/*TM6B, GMR-EYFP, Sb* and *pnr*^*VX6*^*/TM3, twi-Gal4, UAS-2xEGFP, Sb, Ser* were derived from the parental strain *pnr*^*VX6*^/*TM6B, Tb, Hu, e* through genetic crossing with appropriate balancers. The homozygous *pnr*^*VX6-/-*^ embryos (lacking GFP expression) from the *pnr*^*VX6*^/*TM6B, GMR-EYFP, Sb* stock were selected for further immunostaining experiments using anti-coracle and anti-innexin-2 antibodies.

### Genetic interaction experiments

For genetic interaction studies, RNAi-mediated knockdown of *innexin-2* was conducted in a *pannier* heterozygous null mutant (*pnr*^*VX6*^) background. Eggs from both the control and experimental crosses were collected over a 2 hr window at 25°C and subsequently allowed to develop at the same temperature for an additional 12 hours to reach embryonic stage 15/16. Embryos of the control (*UAS-Inx2RNA*i/+; *pnr*^*VX6*^/+) and experimental (*UAS-Inx2RNAi*/+; *pnr*^*VX6*^/*69B-Gal4*) genotypes were then stained using anti-innexin-2, anti-coracle, and anti-spectrin antibodies. In parallel, to serve as an additional control, *innexin-2* knockdown was also performed in a wild-type background (i.e., without the *pannier* heterozygous mutation), following identical egg collection and embryo processing conditions. Embryos of the genotypes *69B-Gal4/+* and *69B-Gal4/UAS-Inx2RNAi* were stained using the same set of antibodies.

### Quantification

For quantifying levels of innexin-2 at the membranes of amnioserosa cells, stacks of confocal images for the control and knockdown embryos were taken and the three slices showing the brightest labelling were used for measurements. Average pixel intensities for 10 junctions per embryo (on 3 confocal slices), 2 embryos of each genotype from two independent experiments, were calculated in Image J using the line measurement tool set to a one-pixel width and background from a laser-off image was subtracted. Knockdown and control mean intensities were averaged and then compared to observe and quantify the changes in expression levels of innexin-2 within the membranes of amnioserosa cells in control and knockdown embryos.

## Results and discussion

### 1. *Pannier* Null Mutants Exhibit Dorsal Closure Defects and Altered Innexin-2 Expression in the Amnioserosa

*Pannier* is expressed in the dorsal most region of *Drosophila* embryos, imagos and adults. As already known, it belongs to the GATA binding class of transcription factors instrumental in defining the dorso-medial region of a fly during development (Calleja *et al*., 2000). Functionally, *pannier* is essential for several developmental processes such as dorsal closure, cuticle formation and specification of the cardiac muscle lineage during fly embryogenesis (Herranz & Morata, 2001; Klinedinst & Bodmer, 2003). In our study involving the *pannier* null mutants, we collected homozygous null mutant embryos of *pannier* from the line of the genotype *pnr*^*VX6*^/*TM6B, GMR-EYFP, S*b. Homozygous null embryos (pnr^*VX6*−/−^) were identified by the absence of GFP and subsequently analysed for dorsal closure defects using immunohistochemistry based approaches. The nulls exhibited incomplete dorsal closure by the end of embryogenesis. Additionally, the amnioserosa cells in the mutants displayed noticeable morphological alterations compared to those in *Canton-S* controls. Although innexin-2 expression remained detectable within the ectoderm, immunostaining revealed a loss of the characteristic amnioserosal staining pattern present in control embryos. This was accompanied by severe alterations in amnioserosal cell morphology and a marked disruption of cell–cell junction integrity, indicating substantial impairment of tissue architecture in the mutants. To determine whether *innexin-2* expression was altered from stage 13 onward, we generated a *pnr*^*VX6*^*/TM3, twi-Gal4, UAS-2xEGFP, Sb, Ser* mutant line and used it for the early identification and staining of embryos. Embryos from this line likewise exhibited disrupted *innexin-2* expression, further supporting the loss of normal gap junction organisation during development in *pannier* null mutants (Supplementary Figure 1). Collectively, these findings validate both the identification of the *pannier* null mutant strain and its suitability for the experiments presented in this study (Figure 2).

**Figure 2:**
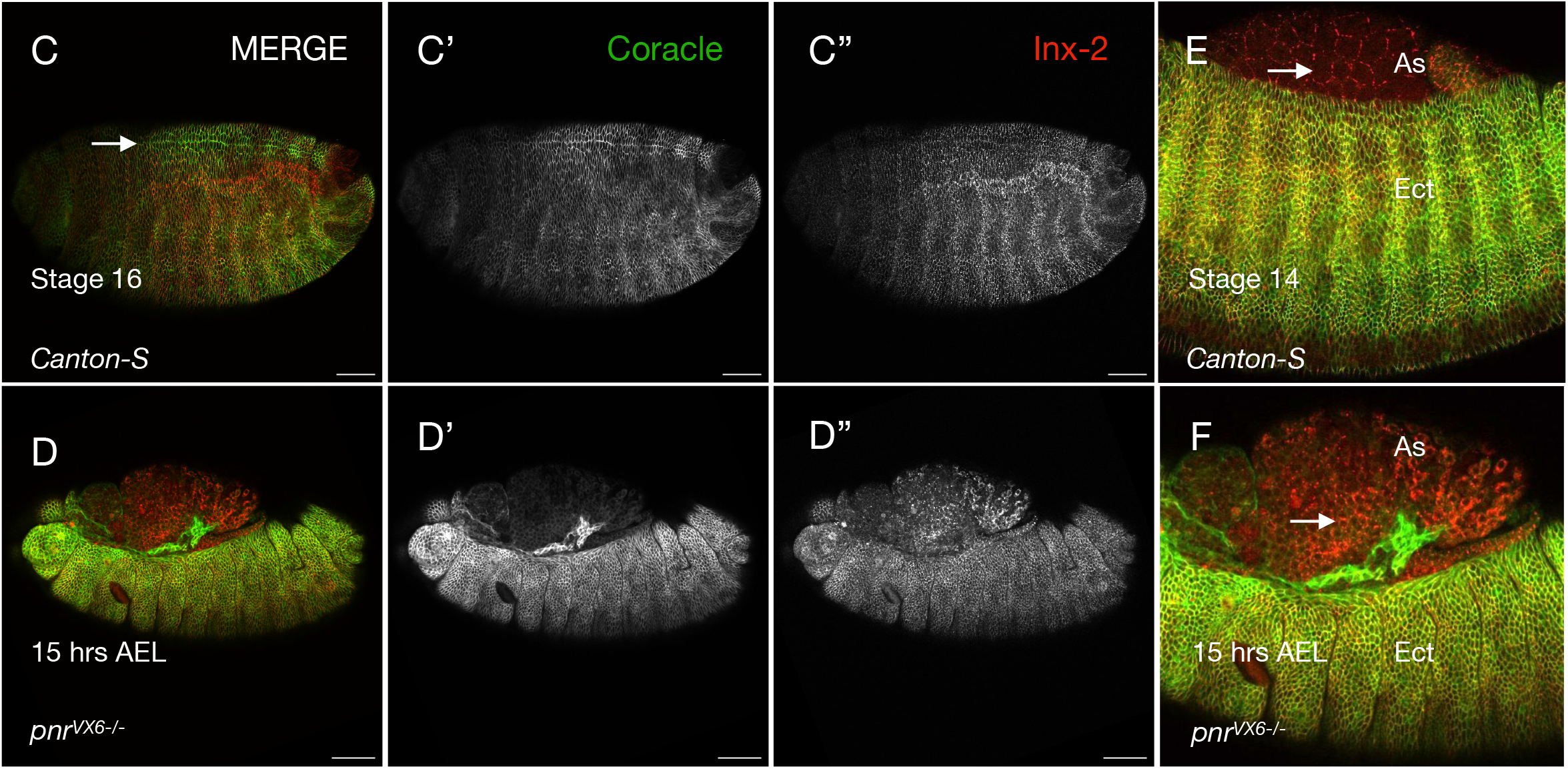
The *pnr*^*VX6-/-*^ mutants exhibit dorsal closure defects, aberrant amnioserosa with disrupted innexin-2 expression. **Panels C-C’’ and D-D’’ :** *Canton-S* and *pnr*^*VX6-/-*^ embryos stained with coracle (green) and innexin-2 (red) antibodies at 15hrs AEL. The dorsal midline seen in *Canton-S* (arrow, C) fail to form in mutants (D) due to closure defects. **E and F :** Magnified views of stage 14 *Canton-S* embryos and dorsally open *pnr* null mutants showing altered amnioserosa morphology and disrupted gap junction expression. In the mutants, the characteristic polygonal shape of amnioserosa cells is not observed. **As** - Amnioserosa, **Ect** - Ectoderm

### 2. Genetic knockdown of *innexin-2*

In fly embryos, during mid embryogenesis, innexin-2 is known to be expressed in the epithelial cells of both the ectoderm and the amnioserosa (Giuliani *et al*, 2013). And by the end of embryogenesis, its expression becomes evident throughout the entire body wall epithelium of the embryo. Additionally, innexin-2 expression is also observed in the epithelial cells of the salivary glands, gut and malphigian tubules (Bauer *et al*., 2005). During embryogenesis, innexin-2 plays essential roles in epithelial morphogenesis (Bauer *et al*., 2004) and foregut development (Lechner *et al*., 2007). The overlap in spatiotemporal expression domains of *pannier* and *innexin-2*, coupled with their established roles in epithelia development and the embryonic lethality observed in their loss-of-function mutants, suggested a potential functional interaction during embryogenesis. Moreover, the involvement of both genes in Decapentaplegic (Dpp) and Wingless (Wg)-dependent developmental processes, albeit in distinct tissue contexts, raised the possibility that they may converge on common regulatory pathways governing embryonic morphogenesis (Zaffran & Tribesman, 2000; Tomoyasu *et al*., 2000; Bauer *et al*., 2002; Bauer *et al*., 2005). To test this possibility, we selectively depleted *innexin-2* using the *pnr-Gal4* and *69B-Gal4* drivers and assessed the resulting developmental phenotypes. RNAi-mediated depletion of *innexin-2* using the *pnr-Gal4* driver resulted in a robust reduction of innexin-2 expression in the cuboidal ectodermal cells. In contrast, the amnioserosa exhibited a heterogeneous response, with embryos displaying different reduced levels of membrane-associated innexin-2, relative to controls (Figure 3, Supplementary Figure 2). The knockdown was accompanied by a specific loss of α-spectrin from the membranes of amnioserosa cells (Figure 3). The observed heterogeneity is consistent with previous observations of *innexin-2* loss-of-function phenotypes and may reflect the persistence of maternally supplied *innexin-2*, which can modulate the penetrance and expressivity of zygotic knockdown effects. Despite these defects, the knockdown embryos completed dorsal closure, however, dorsal closure delays due to affected embryonic development were observed in a few cases. To further assess the requirement of innexin-2 in the embryonic epithelium, we performed RNAi-mediated knockdown using the epithelial-specific *69B-Gal4* driver. This resulted in a marked reduction of innexin-2 expression within the embryonic body-wall epithelium; however, no apparent delay in dorsal closure was observed under these conditions (Figure 4).

**Figure 3:**
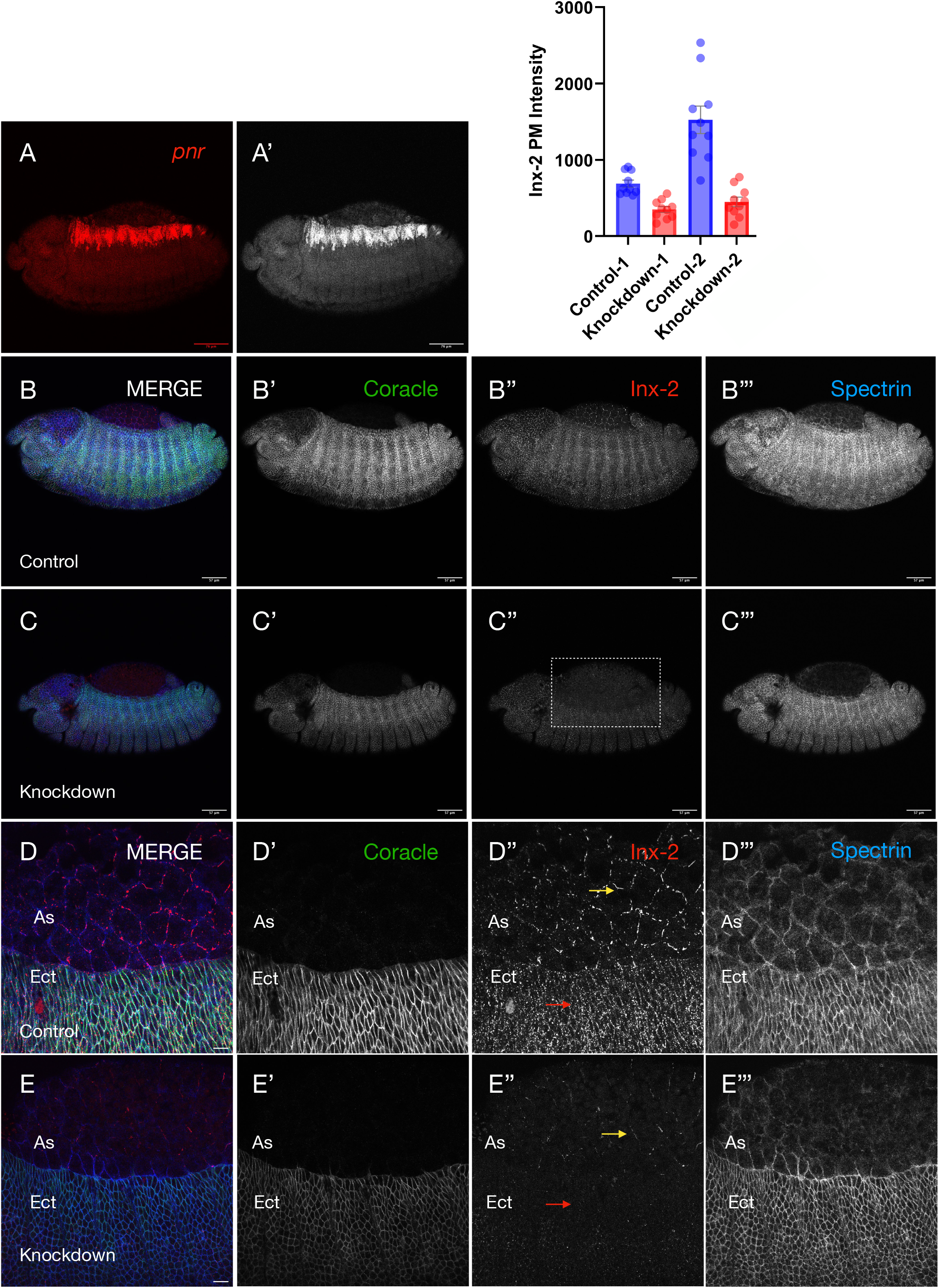
Genetic knockdown of *innexin-2* using *pnr-Gal4* driver. **Panel AA’ :** Stage 13 embryos of genotype *pnr-Gal4/UAS-mCD8-10xRFP* immunostained with anti-RFP (red) to visualise pannier expression. (A) Merged fluorescence image. (A’) Corresponding grayscale image. **Panels B-B’’’ and C-C’’’ :** Control (*pnr-Gal4/+*) and *innexin-2* knockdown (*pnr-Gal4/UAS-wizinx2*) embryos stained with coracle (green), innexin-2 (red), and spectrin (blue), showing altered innexin-2 expression within the amnioserosa (AS) and dorsal epidermis (Ect) (boxed region), along with changes in embryo size. Individual grayscale channels are shown in B’-B’’’ and C’-C’’’. **Panels D-D’’’ and E-E’’’ :** Magnified views of control and knockdown embryos showing the affected ectoderm (Ect) (red arrows in D’’ and E’’) and amnioserosa (As) cells (yellow arrows in D’’ and E’’). **F :** Graphical representation of changes in junctional immunofluorescence levels upon genetic knockdown of *innexin-2* within the amnioserosa cells (n=2 from two independent experiments). Embryos are shown in lateral orientation, with anterior to the left and posterior to the right.

**Figure 4:**
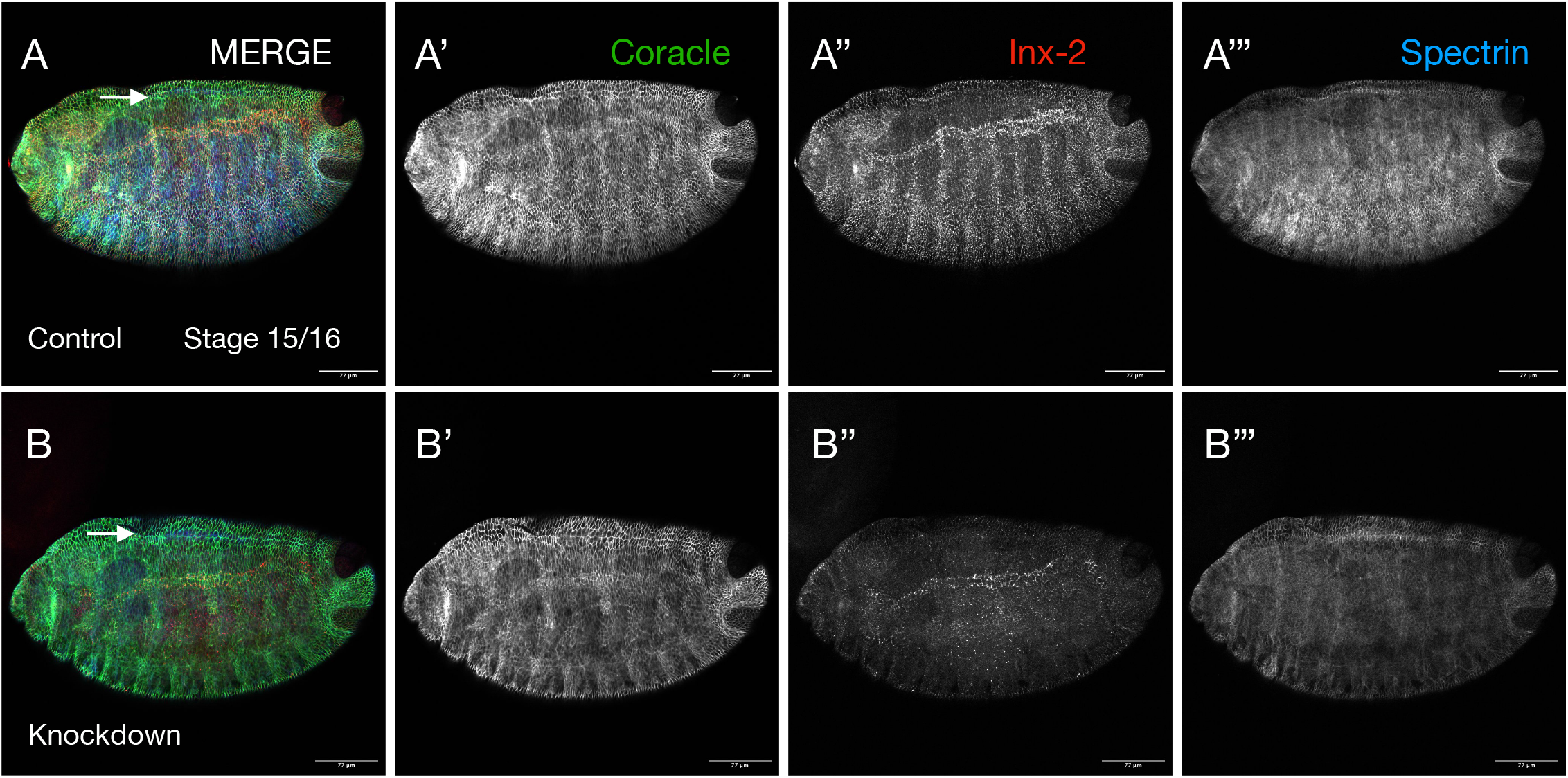
Epidermal knockdown of *innexin-2* utilising *69B-Gal4* driver reduces innexin-2 (red in A’’ and B’’) expression in the embryonic epidermis without affecting dorsal closure by the end of embryogenesis. **Panels A and B :** Composite images of stage 16 control (*69B-Gal4/+*) and knockdown (*69B-Gal4/UAS-Inx2RNAi*) embryos stained with anti-coracle (green), anti-innexin-2 (red), and anti-spectrin (blue) antibodies; **Panels A’–A’’’ and B’–B’’’:** grayscale channels. White arrow in **A** and **B** shows the dorsal midline. Embryos are arranged in a lateral orientation with anterior towards left and posterior towards right.

### 3. Genetic interaction experiments

Genetic experiments in fly have contributed towards understanding the biological functions of innexins in various developmental contexts and processes such as epithelial morphogenesis and regulation of dorsal closure in embryos, development of the embryonic foregut and the central nervous system, modulation of cuboidal to squamous shape transition of the anterior follicle cells (AFCs) in *Drosophila* ovaries and in cell type specification during *Drosophila* oogenesis, regulation of eye disc cell proliferation, morphogenetic furrow movement and amount of differentiated photoreceptors during eye development in flies, crucially required in glial cells for normal post embryonic development of the central nervous system. (Bauer *et al*., 2004; Giuliani *et al*., 2013; Bauer *et al*., 2002; Sahu *et al*., 2017; Sahu *et al*., 2021; Dahl & Muller, 2014, Holcroft *et al*., 2013). Experiments conducted on rats and mice have established that *GATA-4*, a homolog of *pannier*, displays specific interaction with putative binding sites present in the presumed promoters of *Cx40* (Linhares *et al*., 2004). In a separate study providing additional support, it was demonstrated that a distal enhancer for the *Cx30*.*2* gene requires transcription factors Tbx5 and GATA-4 for proper expression in the atrioventricular (AV) conduction system (Munshi *et al*., 2009). The expression pattern of individual connexin show cell-type specificity and developmental changes, indicating distinct but tight control mechanisms for regulation of their expression. Through this study we seek to decipher genetic interaction between *innexins* and *pannier* towards understanding novel functions using the fly embryo as a model system. Genetic interaction studies provide a powerful and useful approach for uncovering functional relationships between genes, enabling the identification of shared regulatory networks, and compensatory mechanisms that collectively govern developmental processes. In *Drosophila*, the availability of sophisticated genetic tools makes such analyses particularly effective for elucidating systems-level mechanisms underlying tissue morphogenesis and pattern formation. To examine the genetic interaction between *innexin-2* and *pannier*, we performed RNAi-mediated knockdown of *innexin-2* using the epithelia-specific *69B-Gal4* driver in a *pannier* heterozygous null background. Experimental embryos (*UAS-inx2RNAi/+; pnr*^*VX6*^*/69B-Gal4*) displayed pronounced defects in body wall epithelial organization, morphology, and size by the end of embryogenesis, compared to control genotypes (*UAS-Inx2RNAi/+; pnr*^*VX6*^*/+* and *UAS-Inx2RNAi/69B-Gal4*) (Figure 5, Figure 6). These findings uncover a previously unrecognized genetic interaction between *pannier* and *innexin-2*, and suggest a novel role for gap junction-mediated communication in the regulation of embryonic size and epithelial morphogenesis (Figure 6). Consistent with this observation, innexins have previously been implicated in the control of cell, tissue, and organ size, volume and growth, including the regulation of embryo size, eye and eye imaginal disc size, and larval brain size (Bauer *et al*., 2004; Richard *et al*., 2017; Richard & Hoch, 2015; Watanabe & Kankel, 1990; Holcroft *et al*., 2013; Das *et al*., 2023; McEvoy *et al*., 2020). Furthermore, innexin-3 colocalizes with innexin-2, exhibits a largely overlapping expression pattern, and has been shown to interact biochemically with innexin-2 at plasma membrane junctions within the embryonic body-wall epithelium (Lehmann *et al*., 2006). Notably, innexin-3 is also required for proper dorsal closure (Giuliani *et al*., 2013), raising the possibility that it contributes to the phenotypes described in the present study or participates in additional morphogenetic processes that were not examined here. Although the roles of innexin-1 and innexin-3 were not directly investigated, their overlapping expression domains and known functional associations with innexin-2 suggest that multiple innexins may act both independently and cooperatively to regulate embryonic development. Future studies combining genetic interaction analyses, tissue-specific knockdown or knockout approaches, and quantitative assessment of epithelial morphogenesis will be important for determining whether innexin-1, innexin-2, and innexin-3 function within a common regulatory network and for elucidating how gap junction-mediated intercellular communication integrates with developmental signaling pathways to coordinate embryonic growth, dorsal closure, and tissue architecture.

**Figure 5:**
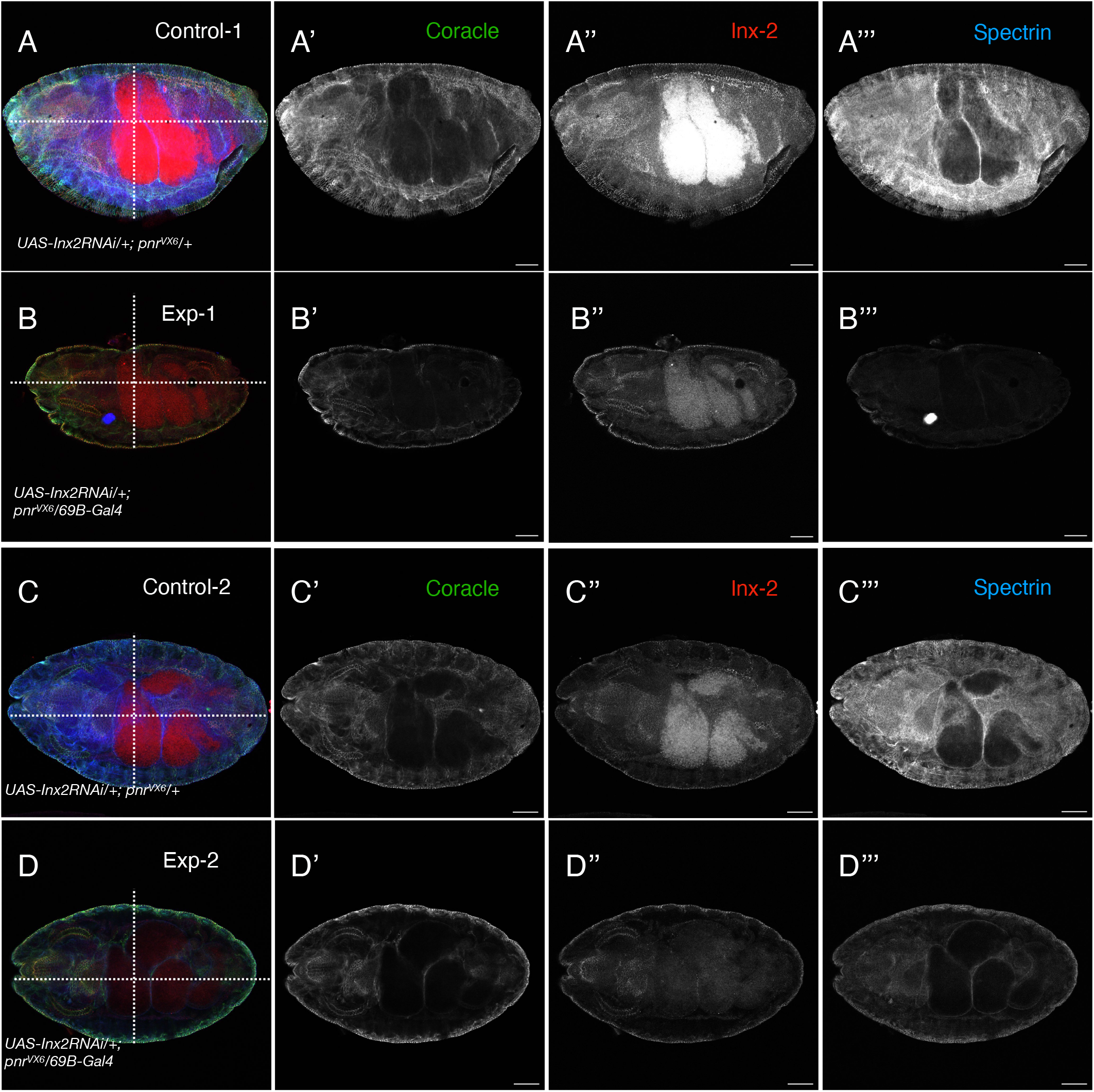
Epidermal knockdown of *innexin-2* using *69B-Gal4* in a *pnr*^*VX6*^ background alters embryo morphology, and results in reduced embryo size by the end of embryogenesis (stage 15/16). **(A–A’’’, C–C’’’) :** Control embryos; **(B–B’’’, D–D’’’) :** Knockdown embryos. Coracle (green), innexin-2 (red), and spectrin (blue) staining are shown as composite and grayscale images. Knockdown embryos **(B**,**D)** exhibit reduced length and width compared to controls **(A**,**C; dotted lines). Panels A and B** show embryos in lateral orientation, while **panels C and D** show embryos in dorsal orientation. In all cases, the anterior end is positioned to the left and the posterior end to the right.

**Figure 6:**
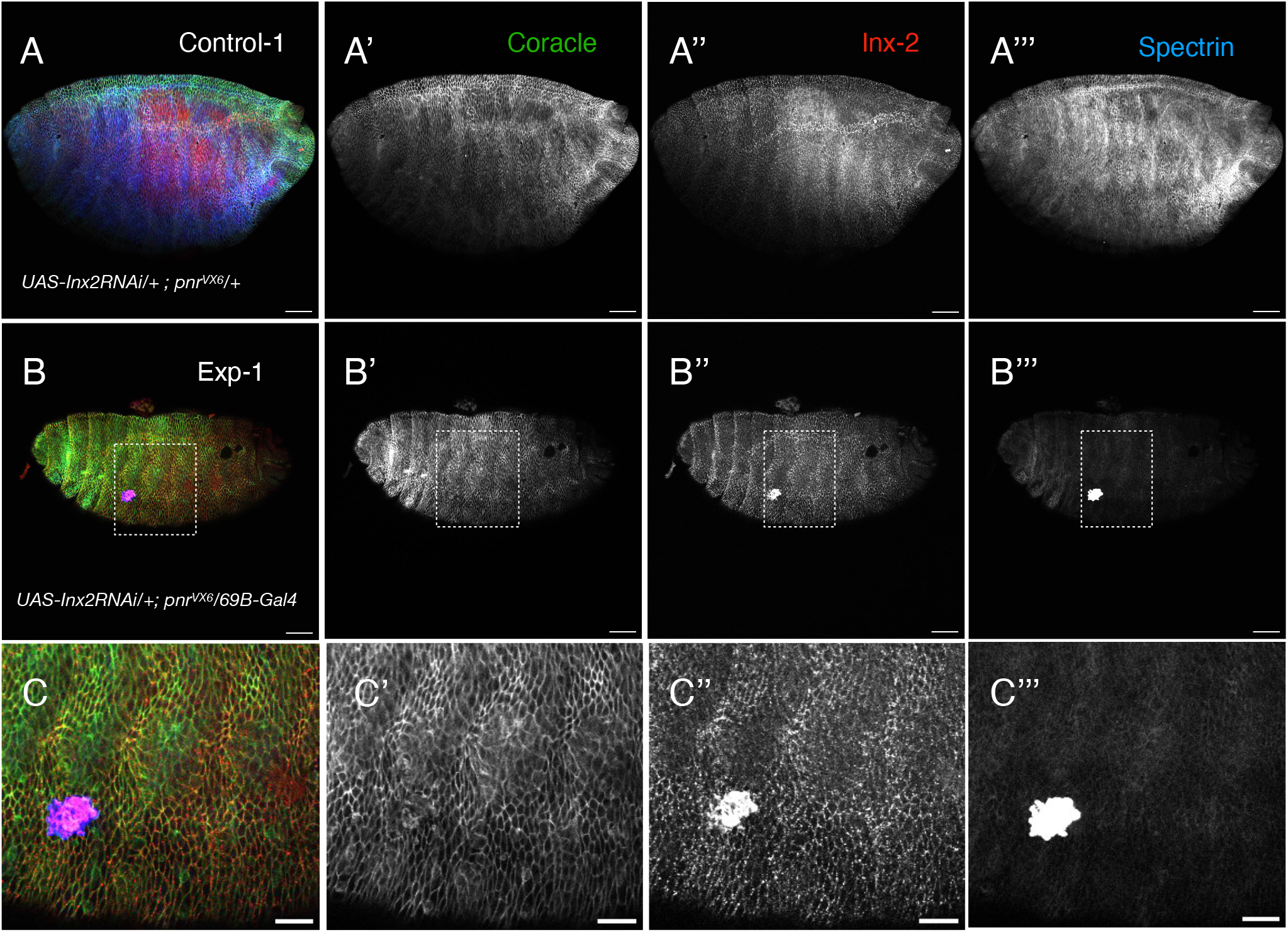
Epidermal knockdown of *innexin-2* using *69B-Gal4* in a *pnr*^*VX6*^ background affects epithelial organisation by late embryogenesis (stage 15/16). **Panels A–A’’’ :** represent control embryos, while **panels B–B’’’ :** depict knockdown embryos. Coracle (green), innexin-2 (red), and spectrin (blue) staining are shown as composite and grayscale images. Knockdown embryos display compromised epithelial integrity, characterized by disrupted cellular architecture and irregular cell packing compared to controls. **Panels C’– C’’’ :** show a magnified view of the boxed region highlighting the altered epithelial morphology in knockdown embryos. Scale bar : 20 µm in panel C-C’’’

## Summary and Outlook

Genetic interaction studies provide a powerful approach for uncovering novel phenotypes and elucidating the functional relationships and regulatory mechanisms governing gap junction proteins and transcription factors. In this study, we identify a previously unrecognized genetic interaction between *pannier* and *innexin-2* that contributes to the regulation of embryonic size and epithelial morphogenesis. The reduction in embryo size observed in the experimental embryos of the genotype *UAS-inx2RNAi/+; pnr*^*VX6*^*/69B-Gal4* compared to controls *UAS-Inx2RNAi/+; pnr*^*VX6*^*/+* and *UAS-Inx2RNAi/69B-Gal4* likely reflects the combined effects of *innexin-2* depletion across a broad range of epithelial tissues targeted by the *69B-Gal4* driver and the compromised amnioserosa cells associated with the *pnr*^*VX6*^ mutation. It will also be important to examine whether embryos carrying combinations of *pnr*^*VX6*^ and *innexin-2* null alleles exhibit comparable defects in embryonic size and morphology, and to assess whether *innexin-2* overexpression can rescue these phenotypes, thereby providing insights into the genetic hierarchy underlying this interaction.

While our findings establish a functional interaction between these genes, the underlying cellular and molecular mechanisms responsible for the observed phenotypes remain to be determined. Furthermore, quantitative approaches such as fluorescence activated cell sorting (FACS)-based analyses may enable more accurate measurements of embryo volume and size differences between genotypes.

In addition to defects in epithelial organisation and embryo size, we observed indications of delayed dorsal closure in the experimental embryos. Although suggestive of developmental delays, these observations require further validation through live-imaging approaches that permit direct assessment of morphogenetic dynamics and developmental timing. Such studies will help distinguish whether the observed phenotypes arise from defects in tissue morphogenesis, altered developmental progression, or a combination of both processes.

## Author contributions

**S.B**.: Conceptualisation, Funding Acquisition, Investigation, Formal Analysis, Data Curation, Visualisation, Writing – Original Draft.

**F.E**.: Funding Acquisition, Training, Writing – Review & Editing.

**R.B**.: Conceptualisation, Funding Acquisition, Resources, Supervision, Writing – Review & Editing.

## Acknowledgements

We sincerely acknowledge the European Molecular Biology Organization (EMBO) for funding support and the NCBS Central Imaging Facility for providing access to imaging infrastructure. We are grateful to the members of AG Bauer at the Hoch/Bauer laboratory, LIMES, University of Bonn, Germany, for their support, inclusivity, participation, valuable scientific inputs, and for generously sharing fly and antibody reagents that contributed to this study. We also thank the Bloomington Drosophila Stock Center for *Drosophila* strains utilised in this work. We gratefully acknowledge the NCBS Fly Facility for fly stock maintenance, husbandry support, and assistance with the management of *Drosophila* resources.

## Declaration

The authors declare no competing interests.

## Use of AI

During the preparation of the manuscript, AI was used to enhance readability and comprehension of the work. The authors subsequently reviewed and edited the content as necessary and take full responsibility for the final version of the publication.

## Notes

### Competing Interest Statement

The authors have declared no competing interest.

