## Supplementary Table 1 for "Genetic interaction approaches reveal emerging roles of innexins in development : Insights from a novel *pannier–innexin-2* interaction during *Drosophila* embryogenesis"

**Supplementary Material**

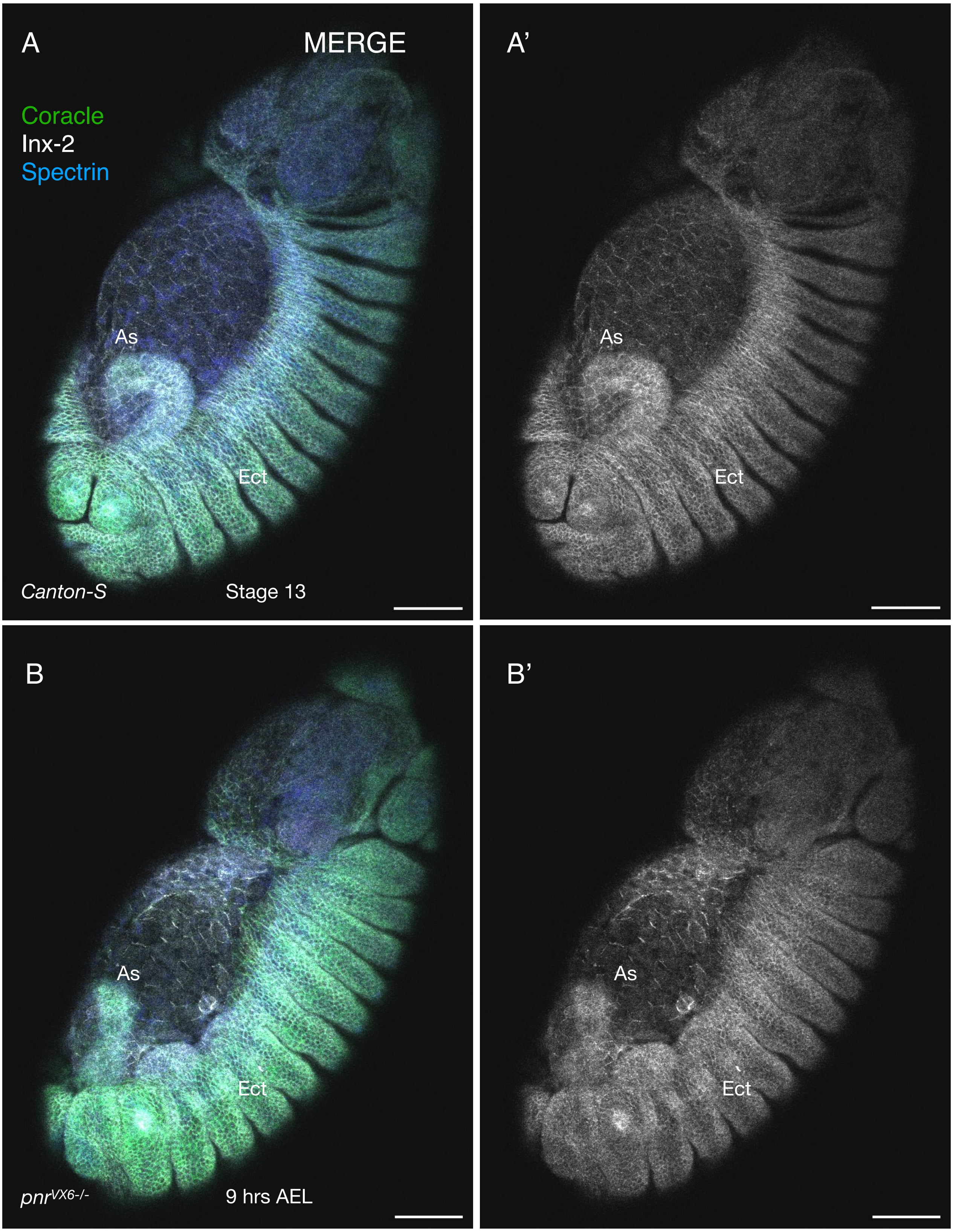


**Supplementary Figure 1 :** The amnioserosa cells of *pannier* null mutants exhibit defects in the levels of innexin-2 expression. **A,A' :** Composite image of a stage 13 embryo showing innexin-2 expression in WT (*Canton-S*) and *pnr*$^{VX6-/-}$ null mutants. **BB' :** Gary scale image of innexin-2 expression in the control and mutant genotypes. As : amnioserosa; Ect : ectoderm.

**Table -1 :** List of fly lines used in this study

| S.No | Genotypes | Source |
|---|---|---|
| 1 | *Canton-S* | BDSC |
| 2 | ; ; *pnr*$^{VX6}$*/TM6B, Tb, Hu, e* | Marc Haenlin, Toulouse, France |
| 3 | ; ; *pnr*$^{VX6}$*/TM6B, GMR-EYFP, S*b | Generated for this study |
| 4 | ; ; *pnr*$^{VX6}$*/TM3, twi-Gal4, UAS-2xEGFP, Sb, Ser* | Generated for this study |
| 5 | ; ; *UASwizInx2* | Hoch/Bauer Lab, Bonn, Germany |
| 6 | ; ; *69B-Gal4* | BDSC 1774 |
| 7 | ; ; *pnr-Gal4/TM6B, Dfd-eYFP, Sb* | Generated for this study |
| 8 | *UAS-Inx2RNA*i/+; *pnr*$^{VX6}$/+ | Generated for this study |
| 9 | *UAS-Inx2RNA*i/+; *pnr*$^{VX6}$*/69B-Gal4* | Generated for this study |
| 10 | ; ; *UAS-Inx2RNAi/69B-Gal4* | Generated for this study |

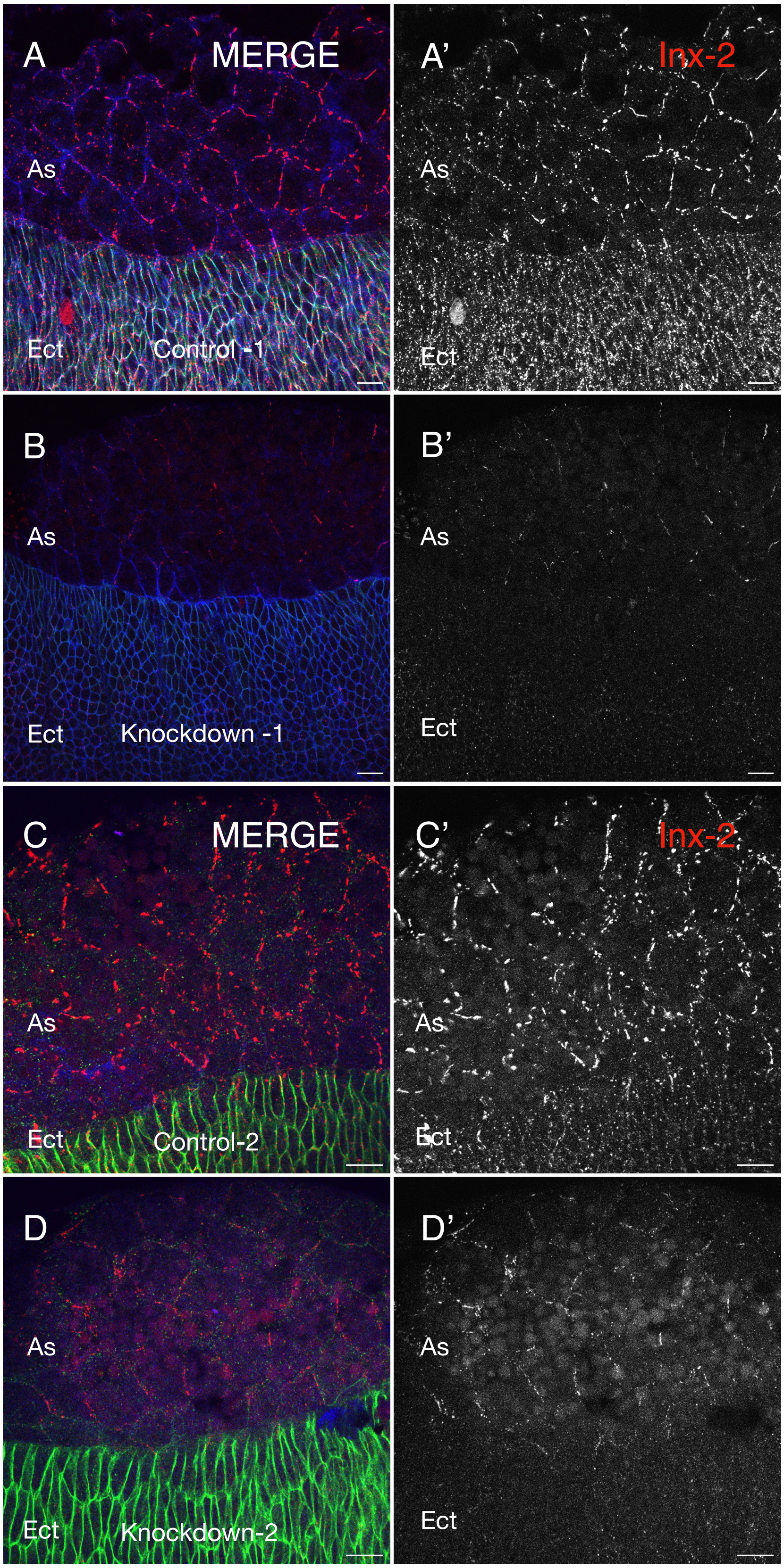


**Supplementary Figure 2 :** Genetic knockdown of *innexin-2* using *pnr-Gal4* driver affects its expression at the membranes of amnioserosa and ectoderm cells. **AA’ , CC’ and BB’ , DD’ :** Merged and gray scale view of control (*pnr-Gal4/+)* and experimental *(pnr-Gal4/UAS-wizinx2*) embryos immunostained using anti-coracle (green), anti-innexin-2 (red) and anti-spectrin (blue) antibodies. As : amnioserosa; Ect : ectoderm
